# The choice of tree prior and molecular clock does not substantially affect phylogenetic inferences of diversification rates

**DOI:** 10.1101/358788

**Authors:** Brice A. J. Sarver, Matthew W. Pennell, Joseph W. Brown, Sara Keeble, Kayla M. Hardwick, Jack Sullivan, Luke J. Harmon

## Abstract

Comparative methods allow researchers to make inferences about evolutionary processes and patterns from phylogenetic trees. In Bayesian phylogenetics, estimating a phylogeny requires specifying priors on parameters characterizing the branching process and rates of substitution among lineages, in addition to others. However, the effect that the selection of these priors has on the inference of comparative parameters has not been thoroughly investigated. Such uncertainty may systematically bias phylogenetic reconstruction and, subsequently, parameter estimation. Here, we focus on the impact of priors in Bayesian phylogenetic inference and evaluate how they affect the estimation of parameters in macroevolutionary models of lineage diversification. Specifically, we use BEAST to simulate trees under combinations of tree priors and molecular clocks, simulate sequence data, estimate trees, and estimate diversification parameters (e.g., speciation rates and extinction rates) from these trees. When substitution rate heterogeneity is large, parameter estimates deviate substantially from those estimated under the simulation conditions when not captured by an appropriate choice of relaxed molecular clock. However, in general, we find that the choice of tree prior and molecular clock has relatively little impact on the estimation of diversification rates insofar as the sequence data are sufficiently informative and substitution rate heterogeneity among lineages is low-to-moderate.

## 3 Introduction

Statistical comparative methods use phylogenetic trees to gain insight into macroevolutionary patterns and processes (Felsenstein 1985; Harvey & Pagel 1991; O’Meara 2012; Rabosky 2016; Harmon 2018). Branch lengths and node ages provide information about the rate of lineage accumulation throughout time (e.g., Nee et al. 1994b; Nee 2006; Ricklefs 2007; Pyron and Burbrink 2013) and are instrumental in characterizing the underlying processes generating global patterns of biodiversity (Schluter & Pennell 2017). A typical workflow uses a point estimate of a phylogenetic tree or a distribution of trees to estimate macroevolutionary parameters, such as the rate of lineage accumulation (speciation) or extinction, which are often compared across groups to provide insight into diversification rates and the tempo of evolution (Nee et al. 1992; Magallón & Sanderson 2001; Alfaro et al. 2009; Rabosky 2014). However, parameter estimates are dependent on the tree from which they are inferred (Felsenstein 1985). Most inference procedures assume that a tree is estimated without error, but, because branch lengths are fundamental to estimates of diversification parameters, uncertain phylogenies can be expected to yield uncertain estimates. A handful of studies have focused on the causes of parameter misestimation when fitting diversification models to trees (e.g., Nee, May, et al. 1994; Barraclough & Nee 2001; Revell et al. 2005; Cusimano & Renner 2010; Rabosky 2010; Moore et al. 2016), but few studies have evaluated uncertainty in phylogenetic estimation explicitly (but see Revell et al. 2005; Wertheim & Sanderson 2011; Marin & Hedges 2018) or characterized aspects of how such uncertainty is quantified, especially in the context of prior specification during Bayesian phylogenetic inference.

Theoretical advances have expanded the scope of phylogenetic comparative methods for studying diversification. Historically, models only assumed a constant rate of lineage diversification or extinction (e.g., Nee et al. 1994b). More modern approaches utilize phylogenies to determine where and/or when shifts in the rates of speciation and extinction take place (see Pyron & Burbrink 2013) or estimate rates that depend on species’ traits (e.g., Maddison et al. 2007; FitzJohn et al. 2009; FitzJohn 2010).

It has been shown that phylogenetic uncertainty and error in tree estimation can directly impact the results of diversification studies. For example, Revell et al. (2005) demonstrated that underparameterization of the model of nucleotide sequence evolution as part of the process of phylogenetic estimation can produce apparent slowdowns in the rate of diversification as quantified by Pybus and Harvey’s (2000) gamma statistic (Revell et al. 2005). Additionally, errors in branch lengths (Wertheim and Sanderson 2011) and biased taxonomic sampling can both affect estimates of diversification rates (Höhna 2014). Taken together, these studies suggest that phylogenetic error can affect the estimation of comparative parameters.

Bayesian methods of phylogenetic inference produce posterior distributions of trees, and comparative parameters can be estimated across such distributions to quantify uncertainty. The use of Bayesian approaches in phylogenetics has increased in recent years due in part to the availability of software, including BEAST (Drummond et al. 2012) and MrBayes (Ronquist et al. 2012). However, the impact that the choice of priors governing the molecular clock and branching process (or “tree prior”) has on the estimate of comparative parameters has not been thoroughly investigated. Two commonly used tree priors are the Yule (Yule 1925) and Birth-Death (BD; Kendall 1948; Nee et al. 1994b; Gernhard 2008; Stadler 2013) models. The Yule model is the simplest of a group of continuous-time branching processes; it has one parameter, *λ*, which is the instantaneous per-lineage rate of speciation that is constant across the tree. The BD model is also a continuous-time process but includes a probability that a lineage will go extinct (and, therefore, leave no descendants); thus, this model has two parameters, *λ* and *µ*, the instantaneous per-lineage rates of speciation and extinction (both of which are constant across the tree). In practice, many approaches re-parameterize the model using *r = (λ − μ*) and *ε = (μ / λ*), the net diversification rate and relative extinction rate, respectively. In general, estimates of *r* have greater precision than *ε* (Nee, May, et al. 1994; Nee, Holmes, et al. 1994; FitzJohn et al. 2009). When using BEAST, researchers must specify a prior distribution on *λ* or on *r* and *ε*, depending on the choice of tree prior.

In addition to priors for branching process parameters, Bayesian phylogenetic analysis also requires the specification of a particular model for rates of evolution across the tree. For example, BEAST gives users the choice of using a strict (or global) molecular clock or an uncorrelated log-normal relaxed molecular clock, among others (Drummond et al. 2012). The strict clock assumes a constant, global rate of sequence evolution across the tree (Zuckerkandl & Pauling 1962), while the uncorrelated log-normal relaxed clock (UCLN) assumes branch-specific rates are drawn from a discretized log-normal distribution independently for every branch in the tree (Drummond et al. 2006). Priors are placed on the mean rate of evolution for the strict clock and the mean and standard deviation of the log-normal distribution for the uncorrelated log-normal relaxed clock.

There is reason to believe that the choice of priors can affect the estimation of diversification parameters. For example, the effects of tree reconstuction on diversification rate estimates were studied by Wertheim and Sanderson (2011). This study focused on trees generated only under a Yule process with a range of *λ* values. The authors simulated sequences under a simple model of sequence evolution (HKY85), and trees were estimated using BEAST assuming a strict clock and narrow prior or range of prior widths on the root age. Their study assessed the impact of sequence length and nodal calibrations on estimating posterior distributions of *λ*, and they found that increasing sequence length leads, as expected, to narrower 95% HPD widths of speciation rates. Additionally, broader calibration priors were shown to increase posterior widths of these estimates. It is plausible that forcing estimation of a tree under a particular branching process (such as a Yule process) may produce an inaccurate tree if the true generating process was different (such as a BD process); this could systematically affect diversification parameter estimates.

Since branch lengths play an important part in estimating diversification parameters, it is also the case that a mismatch of clock models could similarly affect results. Whereas previous work describes a relationship between parameter estimation and misspecification of the model of nucleotide sequence evolution during phylogenetic estimation (Revell et al. 2005), as well as sequence length and nodal calibrations (Wertheim & Sanderson 2011), no studies to our knowledge have directly focused on the impact of tree priors and choice of molecular clocks combined (but see Condamine et al. 2015 for comparisons among Yule and BD priors using an empirical dataset). Additionally, a recent study by Duchěne et al. (2017) emphasizes the importance of appropriately accommodating among-lineage molecular rate variation when inferring diversification rates, both of which may be correlated through underlying evolutionary processes. This study simulated datasets under a variety of diversification conditions with a constant background extinction rate and stressed the importance of accurately capturing variable substitution rates as part of reconstructing the phylogeny.

Here, we quantify the effect of tree prior and clock misspecification on parameter estimation for diversification models. To accomplish this, we simulate phylogenetic trees and associated sequence data under a range of combinations of tree priors and molecular clock models. We then re-estimate trees and use these reconstructed trees to calculate maximum likelihood estimates of diversification rate parameters. We compare these estimates to ones from the original trees to evaluate whether or not priors and clock models contribute to error in estimating diversification rates.

## Materials And Methods

We take advantage of existing applications to simulate trees under a variety of conditions, simulate nucleotide sequence data on these trees, estimate a tree from the nucleotide data, and estimate comparative parameters. The workflow is illustrated in Figure 1. All scripts are written in the R programming language (R Core Team 2015) and are available on GitHub (https://github.com/bricesarver/prior_simulation_study).

**Figure 1.**
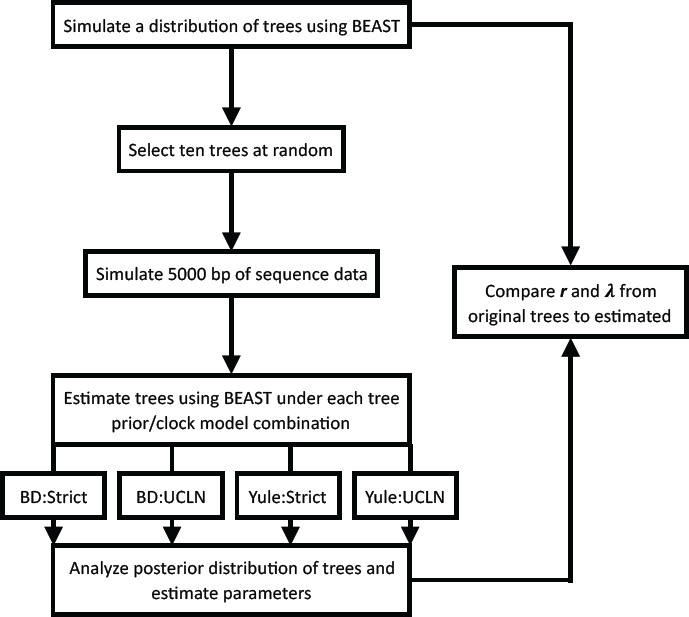
Simulation workflow. *λ* is the instantaneous speciation rate, and *r* is the net diversification rate. Both are estimated for each set of simulation conditions.

### Generation of initial distributions of trees

We simulated trees of two sizes, 25 and 100 taxa, both with a tree depth of 5 arbitrary time units. We simulated initial trees using BEAST v1.7.5 with XML input files generated using BEAUti v1.7.5 (Drummond et al. 2012). DNA sequence data were simulated using these trees with SeqGen v1.3.2 (Rambaut & Grassly 1997).

The simulation process itself consisted of two steps. First, a tree prior was selected for each round of simulations, either Yule or BD. In order to avoid improbable combinations of parameters such that tree shapes were non-randomly sampled (Pennell et al. 2012), initial parameter values for N_t_ were fixed and r calculated using the expectation relating the net diversification rate, the number of taxa, and the tree height: *E[N_t_] = N_0_e^rt^*, where *N_t_* is the number of taxa at *t*, *N_0_* is the initial number of taxa (2 in this case), *r* is the net diversification rate (*λ − μ*), and t is the height of the tree (Nee 2006). Therefore, when N_t_ = 25, *r* = 0.5051, and when N_t_ = 100, *r* = 0.7824, both with a tree height of 5. For BD cases, *ε* was fixed at 0.5.

BEAST requires the specification of a type of molecular clock. For the strict case, the prior on the clock rate was fixed to a log-normal distribution with a mean of 0.01 and a standard deviation of 0.5. For the UCLN case, the prior on the mean of the distribution was of the form U(0.0050, 0.015), and the prior on the standard deviation of the distribution was set to either U(0.17, 0.18), U(0.25, 1), or U(0.25, 1.75). Together, these simulations correspond to a low, medium, and high amount of among-lineage substitution rate heterogeneity.

We then generated a distribution of trees under these conditions using BEAST, sampling only from the priors. To ‘fix’ a parameter, such as root height, to a given value, a normal prior was used with a mean equal to the value and a standard deviation of 0.00001. This prevented BEAST failures using a prior with hard boundary conditions.

### Simulation of nucleotide datasets

For each set of parameter values, we generated a posterior distribution of 10,001 phylograms by sampling from the prior. Ten trees were selected at random without replacement. 5000 bp of sequence data (see Wertheim and Sanderson 2011) were simulated under a GTR+Γ model of nucleotide sequence evolution with parameters estimated in Weisrock et al. (2005) for nuclear rRNA (π_A_: 0.1978, π_C_: 0.2874, π_G_: 0.3403, π_T_: 0.1835; r_AC_: 1.6493, r_AG_: 2.9172, r_AT_: 0.3969, r_CG_: 0.9164 r_CT_: 8.4170, r_GT_: 1.0; α: 2.3592). Sequences were simulated using Seq-Gen v1.3.5 (Rambaut & Grassly 1997) with randomly generated seeds. Additionally, we simulated datasets of two additional sizes, 2500bp and 10000bp, for the 100-taxa, BD:UCLN case to assess the impact of sequence length on parameter estimates. We expect the accuracy of parameter estimates to improve as the amount of sequence data increases owing to more accurate estimation of branch lengths.

### Estimation under tree prior and clock combinations

The resulting NEXUS data files were processed using BEASTifier v1.0 (Brown 2014). BEASTifier takes a list of NEXUS files and generates BEAST XML input files under conditions specified in a configuration file. Each combination of tree priors and clock types was used for each dataset. For example, the sequences generated using a 100 taxon tree that is simulated under a Yule tree prior and strict molecular clock ultimately produced four XML files for analysis: the condition matching the simulation conditions [e.g., a posterior distribution of trees using a Yule tree prior and a strict clock (1)] and all mismatch conditions [e.g., a posterior distribution of trees using a Yule tree prior and a UCLN clock (2), a BD tree prior and a strict clock (3), and a BD prior and UCLN clock (4)]. Each file was then processed using BEAST v1.7.5 (Drummond et al. 2012). Chains were run for 25,000,000 generations (standard analyses) or 50,000,000 generations (additional clock and data-size analyses), sampling every 2500 or 5000, respectively. 10% of the samples (corresponding to 1000 sampled trees) were excluded before analysis as a burn-in. A maximum clade credibility tree was generated for each analysis using TreeAnnotator v1.7.5 assuming median node heights and a posterior probability limit of 0.5.

### Analysis of posterior distributions and maximum clade credibility trees

We analyzed each combination of the four possible simulation/estimation cases (Yule:Strict, Yule:UCLN, BD:Strict, and BD:UCLN) and number of taxa (25 or 100). First, each distribution of trees was rescaled to the exact root height of the original tree using ape (Paradis et al. 2004). This was performed to remove any confounding effects that may have been introduced when the estimating the root age. Then, for each tree in the posterior, we estimated *λ* and *r* by maximum likelihood using the DDD package in R (Etienne et al. 2012; Etienne & Haegeman 2012).

In addition, we produced lineage-through-time (LTT) plots for each replicate. The LTT plot of the maximum clade credibility tree produced from each analysis was plotted on the same graph as the original tree from which the data were simulated. Each plot, then, consists of LTT plots for the 10 original trees and consensus trees from the corresponding 10 posterior distributions.

## Results and Discussion

The goal of this study is to determine the impact the choice of tree prior and molecular clock have on the estimation of comparative phylogenetic parameters. We focused our efforts on estimating *λ*, the rate of lineage accumulation, and *r*, the net diversification rate, under all combinations of two tree priors (Yule and BD) and two flavors of molecular clocks (strict and UCLN). These parameters were selected for investigation because estimating the relative extinction rate (*ε*) alone is known to be difficult, and estimates of this parameter have larger uncertainty (e.g., Nee, Holmes, et al. 1994). Estimating the net diversification rate still provides insight into the effect of extinction across the phylogeny while facilitating a meaningful comparison among simulation conditions. We found that the combination of tree prior and clock did not substantially impact diversification parameter estimates. Across our simulation conditions, parameters from trees estimated under all combinations of tree priors and clocks were concordant with parameter estimates produced from the trees on which nucleotide data were simulated. When original trees were simulated under a Yule process, all combinations of tree priors and clocks produced extremely similar estimates to the parameters estimated from trees on which data were simulated (Figure 2). Distributions overlapped across all combinations of tree priors and molecular clocks. Slight deviations from simulated values are likely attributable to sampling error. The estimates of *λ* and *r* were consistently underestimated for the 25-taxa UCLN cases, providing evidence that the number of taxa is important when among-lineage rate heterogeneity is concerned. However, other preliminary trials did not show a consistent pattern of underestimation, suggesting that this pattern results from the 10 trees initially selected for simulation and not a systematic bias (data not shown). LTT plots of maximum clade credibility trees indicated that the estimated trees generally coincide with the original trees, though the Yule:UCLN case showed greater discordance at nodes deeper in the tree for a small number of replicates (Supplementary Figure S1). This is not surprising given the difficulty of estimating nodes deep in the tree, and it also helps explain the discrepancy described above.

**Figure 2.**
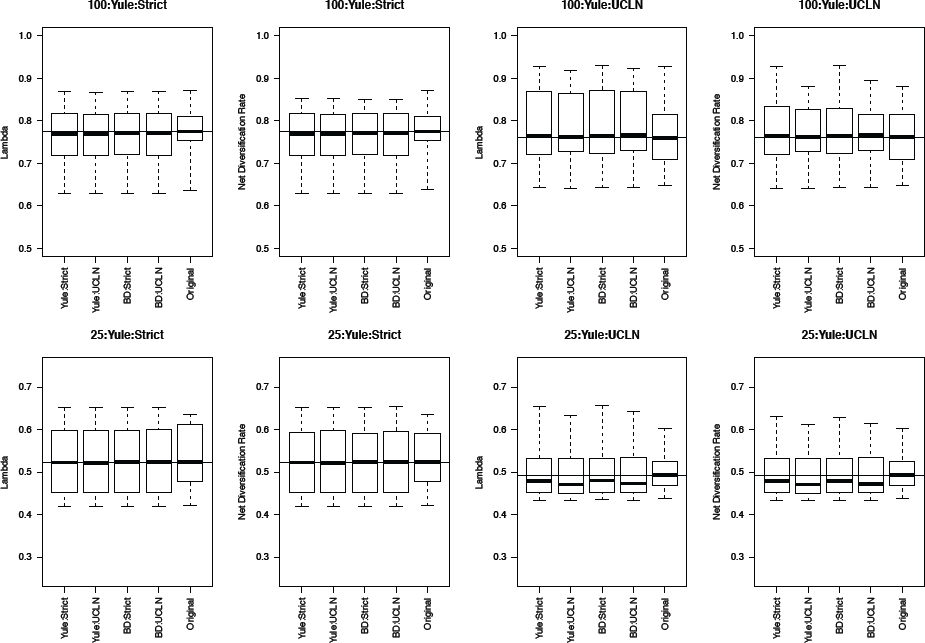
Yule simulations. The top row of plots refers to the 100-taxa cases, whereas the bottom row refers to the 25-taxa cases. Median estimates of *λ* or *r*, estimated from the 10 original trees, are used as data for each boxplot. The title of each subplot refers to the simulation conditions. Each combination of tree priors and molecular clocks under which trees are estimated is listed on the x-axis. The distribution of estimates from the original trees is also displayed. Parameter estimates are generally consistent with the original trees with slight deviations in some cases.

When trees were simulated under a BD process, estimates were also generally concordant with the original trees. Medians were nearly identical among many simulation conditions (Figure 4), though parameters were underestimated in the UCLN cases. This discrepancy was either reduced or did not appear to be present in cases assuming a strict clock. LTT plots revealed that maximum clade credibility trees were, again, approximately equivalent to the original. There were some exceptions, again in the deep nodes of the trees, though these did not drastically affect parameter estimation (Figure 3). As in the Yule cases, there were no discernable tendencies for parameter estimates to be consistently over or underestimated relative to the simulated trees in preliminary analyses (data not shown). However, estimates of *λ* are biased downwards, sometimes drastically. For the 25-taxa cases, *λ* estimates are close to *r*, even though they ought to be 2*r* with *ε* = 0.5. We hypothesize that estimates of *λ* should approach 2*r* as the number of taxa increases. To investigate, we performed additional simulations, as described above, but with 50, 75, and 125 taxa. Estimates of lambda increase with the number of taxa but are still reduced (Supplementary Figure S3). This suggests that estimates of *λ* may be incorrect when trees are estimated assuming no extinction.

**Figure 3.**
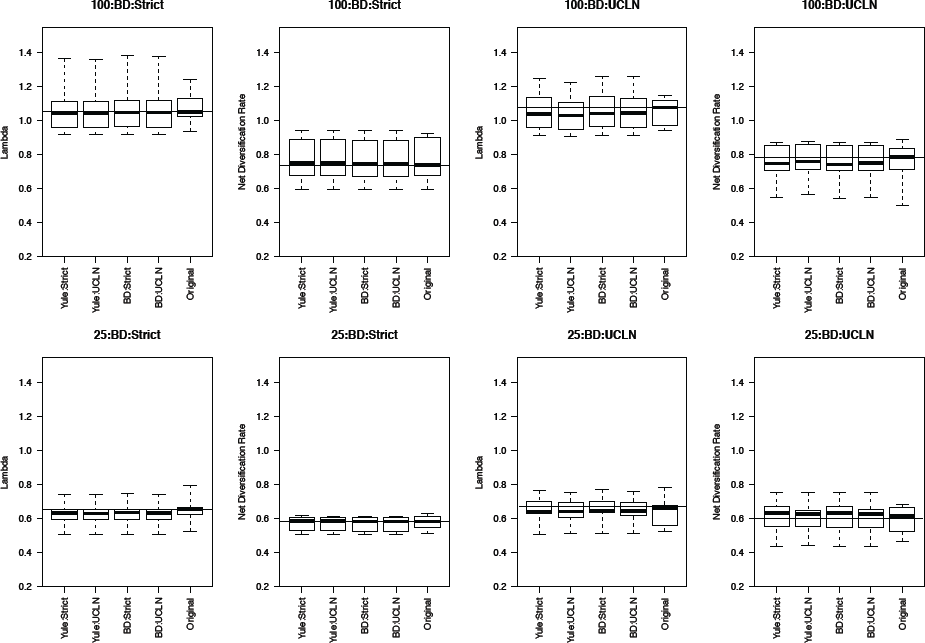
Birth-Death simulations. The top row of plots refers to the 100-taxa cases, whereas the bottom row refers to the 25-taxa cases. The median estimates of *λ* or *r*, estimated from the 10 original trees, are used as data for each boxplot. The title of each subplot refers to the simulation conditions. Each combination of tree priors and molecular clocks under which trees are estimated is listed on the x-axis. The distribution of estimates from the original trees is also displayed. Parameter estimates are highly congruent with the original trees under each set of simulation conditions.

**Figure 4.**
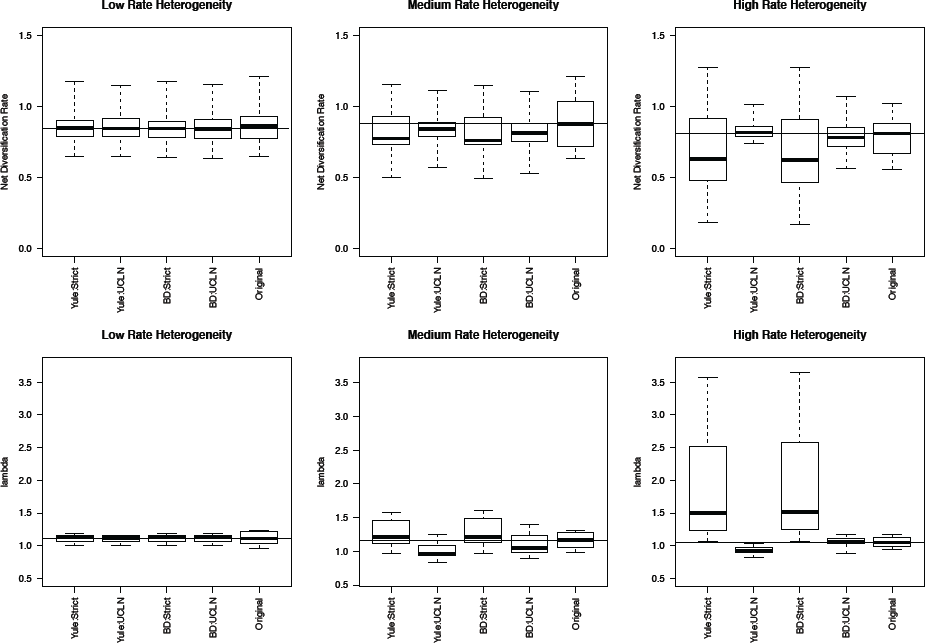
Effect of substitution rate heterogeneity on diversification rate estimates. Three simulated datasets with low, medium, and high substitution rate heterogeneity (see Materials and Methods) are displayed. Parameter estimates agree with simulated data in the low heterogeneity case across all combinations of priors. The deviation of estimates from the original trees increases as the amount of heterogeneity increases. The effect is most pronounced in the ‘high’ case, where the use of a strict molecular clock fails to capture heterogeneity and produces substantially different estimates. An uncorrelated log-normal molecular clock produces reasonable estimates in all cases.

The simulations involving low, medium, and high among-lineage substitution rate heterogeneity revealed that it is possible for the choice of clock to have a substantial impact on parameter estimates (Figure 4). With low rate heterogeneity, estimates of *λ* and *r* are similar to the original trees, but the discordance increases dramatically as the variance in rates among lineages increases. Trees estimated using an uncorrelated lognormal clock appear to suffer the least, especially when estimated under the simulation conditions (BD:UCLN). This effect is most dramatic in the high rate heterogeneity simulations, where the assumption of a tree-wide constant substitution rate can lead to substantially discordant estimates of both *λ* and *r*. Furthermore, it is worth noting that analysis of each of these simulation conditions indicates a deviation from a strict clock, as evidenced by posterior estimates of the coefficient of variation from BEAST on the simulated datasets (95% HPD, low rate hetereogeneity: [0.157, 0.1948]; medium rate heterogeneity: [0.2508, 1.1833]; high rate heterogeneity: [0.2402, 2.6596]). In other words, investigators could easily avoid errors associated with using a strict clock by testing for rate heterogeneity in their sequence data.

In conclusion, it appears that reasonable parameter estimates can often be achieved with either prior. Among these simulated cases, either choice of tree prior appears to capture the underlying branching process on which data was simulated; the same holds for molecular clocks with low among-lineage rate heterogeneity. While estimates are concordant across tree priors and clock models, previous studies have shown that the accuracy of the estimates depends on the amount of data available; here, this refers to the number of taxa. In one example, trees of 1000 taxa produce more accurate estimates of diversification rates than trees of 100 taxa (Stadler 2013). Our results are consistent with Stadler’s conclusions. Adding more informative data produces more accurate phylogenetic estimates (assuming no signal conflict) and should reduce the impact of stochasticity on parameter estimation.

The assumption of a single rate of evolution across a tree is often violated and can severely impair phylogenetic estimation (e.g., Shavit et al. 2007; Penny 2013). This study assumed rates with a modest amount of heterogeneity, and it appears that a strict clock produces reasonable results in the face of this violation. In other words, a dataset with a small to moderate amount of heterogeneity may have rates that are reasonably captured by a single, global rate. However, it may not be known *a priori* whether a dataset has disparate rates of evolution among lineages. It would be advisable, then, to assume a clock model that has the potential to model heterogeneity more accurately, and this is partially why the uncorrelated log-normal relaxed clock has seen such widespread use and success in systematic analyses (Drummond et al. 2006). Furthermore, should rates of evolution be extreme among some lineages, it would make sense to attempt to capture any heterogeneity using appropriate priors as opposed to assuming it is absent. Rate homogeneity among lineages, or the absence of a clock altogether, may represent a poor prior given our current understanding of molecular biological processes (Drummond et al. 2006).

There are several caveats to these conclusions. First, our original trees are fully resolved, and nucleotide sequence data are simulated under parameters estimated from a quickly evolving nuclear intron. This indicates that there will be a large number of phylogenetically informative sites per individual. Therefore, these trees will be easier to estimate than those that lack signal and/or contain unresolved nodes. Second, there is no extreme rate heterogeneity among lineages. Third, the datasets only contain 25 and 100 taxa, each with only 5000 bp of nucleotide sequence data, following the protocol of Wertheim and Sanderson (2011). Datasets of this size are considered modest in the current era of high-throughput sequencing, where the generation of hundreds of thousands or millions of base pairs of sequence per individual is possible. More sequence data can lead to more accurate phylogenies, which improves parameter estimates at the expense of computational speed. It is also reasonable to assume that some systems may be best explained through more complex models, i.e., models that specifically assume multiple, independent diversification rates across a dataset (e.g., Alfaro et al. 2009; Rabosky 2014). Our analyses only assume a single rate of diversification, and this assumption may be violated in larger datasets with greater levels of taxonomic divergence. It is important to select among models in order to produce accurate, interpretable results for each dataset.

## 15 Funding

This work was supported by the National Science Foundation (DEB-0717426 to B.A.J.S. and J.S.; DEB-1208912 to L.J.H.). B.A.J.S., J.S. and K.M.H. received funding from BEACON, a National Science Foundation-funded Center for the Study of Evolution in Action (DBI-0939454). Additionally, M.W.P. was supported by a National Sciences and Engineering Research Council of Canada post-graduate fellowship. This project used the IBEST Computational Resources Core, supported by grants from the National Center for Research Resources (5 P20 RR016448-10) and the National Institute of General Medical Sciences (8 P20 GM103397-10) from the National Institutes of Health. Any opinions, findings, and conclusions or recommendations expressed in this material are those of the authors and do not necessarily reflect the views of the National Science Foundation.

### Acknowledgements

The authors thank Jonathan Eastman for helpful discussion on earlier versions of this project and Rob Lyon for expert help through the IBEST Computational Resources Core. We also thank Frank Burbrink for helpful comments and insights on an earlier version of this manuscript.

---

**Supplementary Figure S1.**
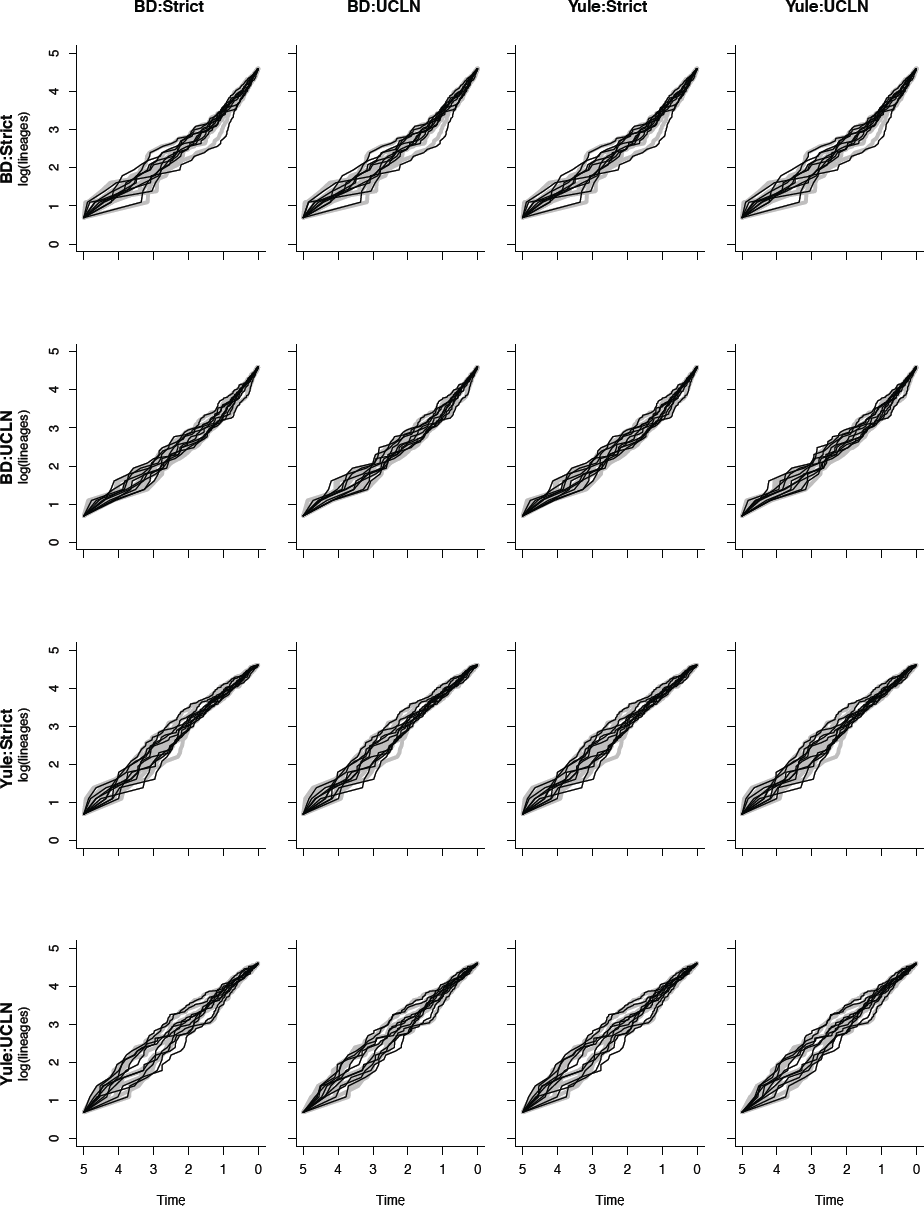
Lineagethroughtime plots. The yaxis of each plot is natural-log transformed. Rows refer to conditions under which original trees are simulated, and columns refer to conditions under which trees are estimated. Thick gray lines represent the original trees and are, therefore, identical across each row of plots. Thin dark lines refer to the maximum clade credibility trees summarized from the posterior distribution of trees under the specified combination of tree prior and molecular clock. There is a significant amount of concordance, indicative of accurate phylogenetic estimation, though some discordance (indicated by nonoverlapping lines) is revealed.

**Supplementary Figure S2.**
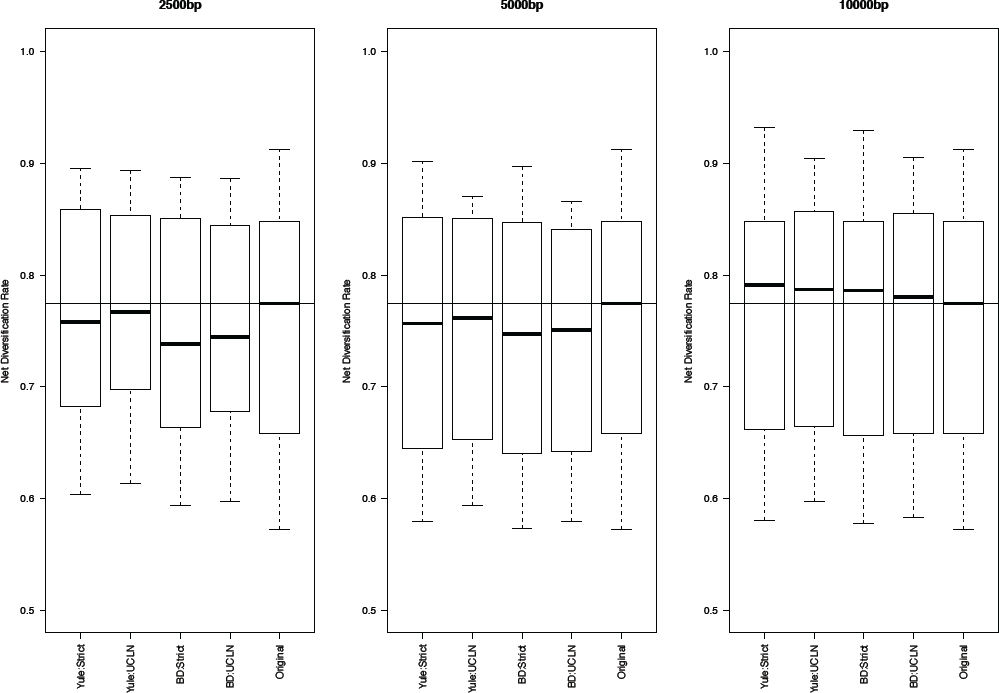
The effect of data size on diversification rate estimates. Estimates of net diversification rate approach the values of the original simulations as the size of the dataset increases, especially for the combination of tree prior and molecular clock under which the data were simulated (i.e., BD:UCLN).

**Supplementary Figure S3.**
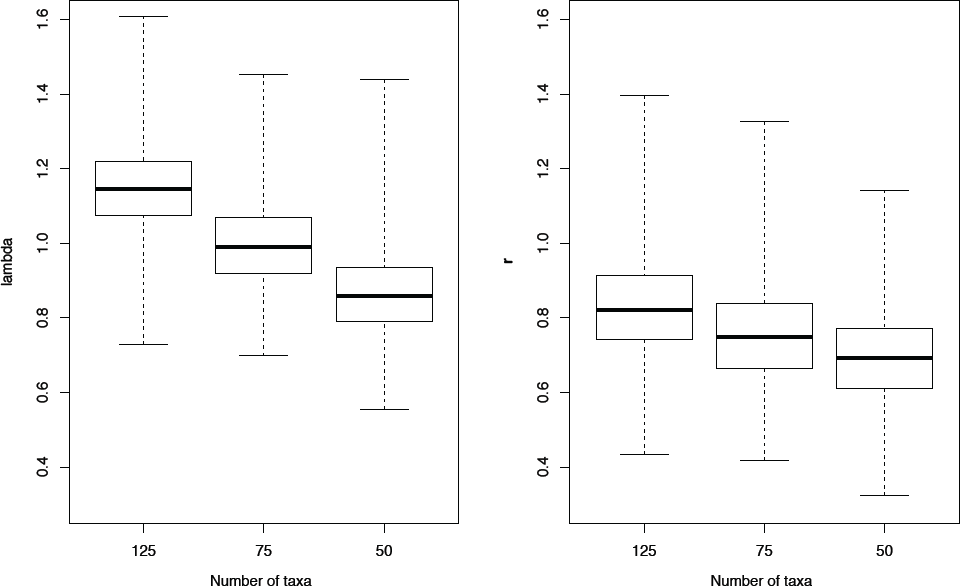
The effect of the number of taxa on estimates of net diversification rate and speciation rate. The xaxis (125, 75, or 50) corresponds to the number of taxa, and the yaxis corresponds to the value of the parameter estimated (*λ* or *r*). Since posterior distributions were generated under a birthdeath process with the relative extinction rate equal to 0.5, estimates of *λ* should approach 2*r*. While the difference from the expectation improves with the number of taxa, the discrepancy can be attributed to a combination of sample size and model misspecification (i.e., estimating *λ* assuming a Yule process when the generating model was BirthDeath).

